# Deep Learning-based Pipeline to Recognize Alzheimer’s Disease using fMRI Data

**DOI:** 10.1101/066910

**Authors:** Saman Sarraf, Ghassem Tofighi

**Affiliations:** Department of Electrical and Computer Engineering, McMaster University, Hamilton, ON, L8S 4L8, Canada, Rotman Research Institue at Baycrest, University of Toronto; Electrical and Computer Engineering Department, Ryerson University, Toronto, ON M5B 2K3 Canada

**Keywords:** Deep learning, Alzheimer’s Disease, fMRI

## Abstract

Over the past decade, machine learning techniques and in particular predictive modeling and pattern recognition in biomedical sciences, from drug delivery systems to medical imaging, have become one of the most important methods of assisting researchers in gaining a deeper understanding of issues in their entirety and solving complex medical problems. Deep learning is a powerful machine learning algorithm in classification that extracts low- to high-level features. In this paper, we employ a convolutional neural network to distinguish an Alzheimers brain from a normal, healthy brain. The importance of classifying this type of medical data lies in its potential to develop a predictive model or system in order to recognize the symptoms of Alzheimers disease when compared with normal subjects and to estimate the stages of the disease. Classification of clinical data for medical conditions such as Alzheimers disease has always been challenging, and the most problematic aspect has always been selecting the strongest discriminative features. Using the Convolutional Neural Network (CNN) and the famous architecture LeNet-5, we successfully classified functional MRI data of Alzheimers subjects from normal controls, where the accuracy of testing data reached 96.85%. This experiment suggests that the shift and scale invariant features extracted by CNN followed by deep learning classification represents the most powerful method of distinguishing clinical data from healthy data in fMRI. This approach also allows for expansion of the methodology to predict more complicated systems.

## I. Introduction

Alzheimers disease is an irreversible, progressive neurological brain disorder. It is a multifaceted disease that slowly destroys brain cells, causing memory and thinking skill losses, and ultimately loss of the ability to carry out even the simplest tasks. The cognitive decline caused by this disorder ultimately leads to dementia. For instance, the disease begins with mild deterioration and gets progressively worse as a neurodegenerative type of dementia. Diagnosing Alzheimers disease requires very careful medical assessments, such as patient history, a mini mental state examination (MMSE), and physical and neurobiological exams. In addition to these evaluations, resting-state functional magnetic resonance imaging (rs-fMRI) provides a non-invasive method of measuring functional brain activity and changes in the brain [13]. There are two important concepts regarding resting-state fMRI. First, because patients do not perform any tasks and there is no simulation, the procedure is more comfortable than a normal fMRI. Second, rs-fMRI data acquisition can be performed during a clinical scan, and most researchers are interested in brain network analysis and extraction from rs-fMRI data [8][9][11][6][2]. However, development of an assistive tool or algorithm to classify fMRI data and, more importantly, to distinguish brain disorder data from healthy subjects has always been of interest to clinicians. Any machine learning algorithm that is able to classify Alzheimers disease will assist scientists and clinicians in diagnosing this brain disorder. In this work, the convolutional neural network (CNN), which is part of the Deep Learning Network architecture, is utilized in order to distinguish Alzheimers brains from healthy brains and produce a trained, predictive model.

## II. Background and Algorithms

### A. Data Acquisition and Preprocessing

In this work, 28 Alzheimer’s disease (AD) patients and 15 normal control (NC) subjects (24 female and 19 male) with a mean age of 74.9 5.7 years were selected from the ADNI^1^ dataset. The AD subjects’MMSEs were reported to be over 20 by ADNI, and normal participants were healthy, with no reported history of medical or neurological conditions. Scanning was performed on a Trio 3 Tesla, which included structural and functional scans. First, anatomical scans were performed with a 3D MP-RAGE sequence (TR=2s, TE=2.63 ms, FOV=25.6 cm, 256 × 256 matrix, 160 slices of 1mm thickness). Next, functional scans were obtained with an EPI sequence (150 volumes, TR=2 s, TE=30 ms, flip angle=70, FOV=20 cm, 64 × 64 matrix, 30 axial slices of 5mm thickness, no gap). The fMRI data were pre-processed using the standard modules from the FMRIB Software Library v5.0 [10]. Preprocessing steps for the anatomical data involved the removal of non-brain tissue from T1 structural images using the Brain Extraction Tool. Preprocessing steps for the functional data included motion correction (MCFLRIT), skull stripping, and spatial smoothing (Gaussian kernel of 5-mm FWHM). Low-level noise was removed using high-pass temporal filtering ($sigma=90.0 sec). Functional images were then aligned to individual high-resolution T1-weighted scans, which were subsequently registered to the Montreal Neurological Institute standard space (MNI152) using affine linear registration and resampled at 2mm cubic voxels. The end results of the preprocessing step were 45x54x45x300 images, from which the first 10 slices of each image were removed, as they contained no functional information.

### B. Deep Learning

Hierarchical or structured deep learning is a modern branch of machine learning that was inspired by the human brain. This technique has been developed based on complicated algorithms that model high-level features and extract those abstractions from data by using neural network architecture that is similar but much more complicated. Neuroscientists have discovered that the neocortex, which is a part of the cerebral cortex concerned with sight and hearing in mammals, processes sensory signals by propagating them through a complex hierarchy over time. This served as the primary motivation for developing deep machine learning that focuses on computational models for information representation which exhibits similar characteristics to those of the neocortex [1][4][3].

#### 1) Convolutional Neural Networks (CNNs / ConvNets)

Convolutional Neural Networks that are inspired by the human visual system are similar to classic neural networks. This architecture has been specifically designed based on the explicit assumption that raw data is comprised of twodimensional images that enable us to encode certain properties and also reduce the amount of hyper parameters. The CNN topology utilizes spatial relationships to reduce the number of parameters that must be learned and thus improves upon general feed-forward back propagation training. Equation 1 demonstrates how Error is calculated in the back propagation step, where E is error function, y is the i^th^, j is the neuron, x is the input, l represents layer numbers, w is the filter weight with a and b indices, N is the number of neurons in a given layer, and m is the filter size.

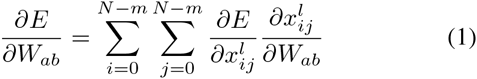

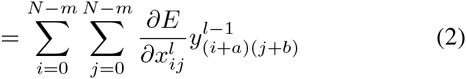

In CNNs, small portions of the image (dubbed local receptive fields) are treated as inputs to the lowest layer of the hierarchical structure. One of the most important features of CNN is that the complex architecture provides a level of invariance to shift, scale and rotation, as the local receptive field allows the neuron or processing unit access to elementary features such as oriented edges or corners. This network is basically constructed of neurons having learnable weights and biases and forming a Convolutional Layer. It also includes other network structures, such as a Pooling Layer, a Normalization Layer, and a Fully-Connected Layer. As briefly mentioned above, the Convolutional Layer, or so-called CONV, computes the output of neurons that are connected to local regions in the input. Each computes a dot product between its weight and the region it is connected to in the input volume. The Pooling Layer, or so-called POOL, performs a down sampling operation along the spatial dimensions. The Normalization Layer, or RELU, applies an element-wise activation function, such as the max (0, x), thresholding at zero. This layer does not change the size of the image volume. The Fully-Connected Layer, or FC, computes class scores, resulting in the volume of the number of classes. As with ordinary Neural Networks, and as the name implies, each neuron in this layer is connected to all of the numbers in the previous volume [1] [4]. The Convolutional Layer plays an important role in CNN architecture and is the core building block in this network. The CONV layers parameters consist of a set of learnable filters. Every filter is spatially small but extends through the full depth of the input volume. During the forward pass, each filter is convolved across the width and height of the input volume, producing a 2D activation map of that filter. During this convolving, the network learns of filters that activate when they see some specific type of feature at some spatial position in the input. Next, these activation maps are stacked for all filters along the depth dimension, which forms the full output volume. Thus, every entry in the output volume can also be interpreted as an output of a neuron which only looks at a small region in the input and shares parameters with neurons in the same activation map [1] [4] [3]. A Pooling Layer is usually inserted between successive Convolutional Layers in ConvNet architecture. Its function is to reduce (down sample) the spatial size of the representation in order to control the amount of network hyper parameters, and hence to also control overfitting. The Pooling Layer operates independently on every depth slice of the input and resizes it spatially using the MAX operation. Recently, more successful CNNs have been developed, such as LeNet, AlexNet, ZF Net, GoogleNet, VGGNet and ResNet. The major bottleneck of constructing ConvNet architecture is the memory restrictions of GPU [1] [4] [3]. As is evident in Figure 3, LeNet-5 was first designed by Y. LeCun et al. [4], and this famous network successfully classified digits and was applied to hand-written check numbers. The application of this network was expanded to more complicated problems, and the hyper parameters were adjusted for new issues. However, more sophisticated versions of LeNet have been successfully tested. In this work, we addressed a very complicated binary classification of Alzheimers data and normal data. In other words, we needed a complicated network for two classes, which compelled us to choose LeNet-5 and adjust this architecture for fMRI data.

The implemented network is shown in Figure 2 in detail.

**Fig. 1:**
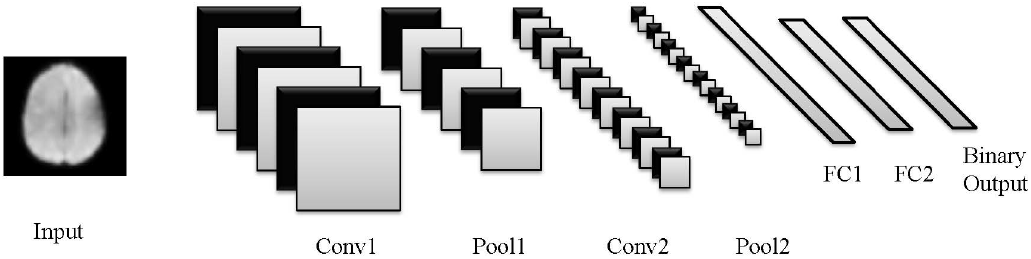
LeNet-5 architecture adopted for fMRI data

**Fig. 2:**
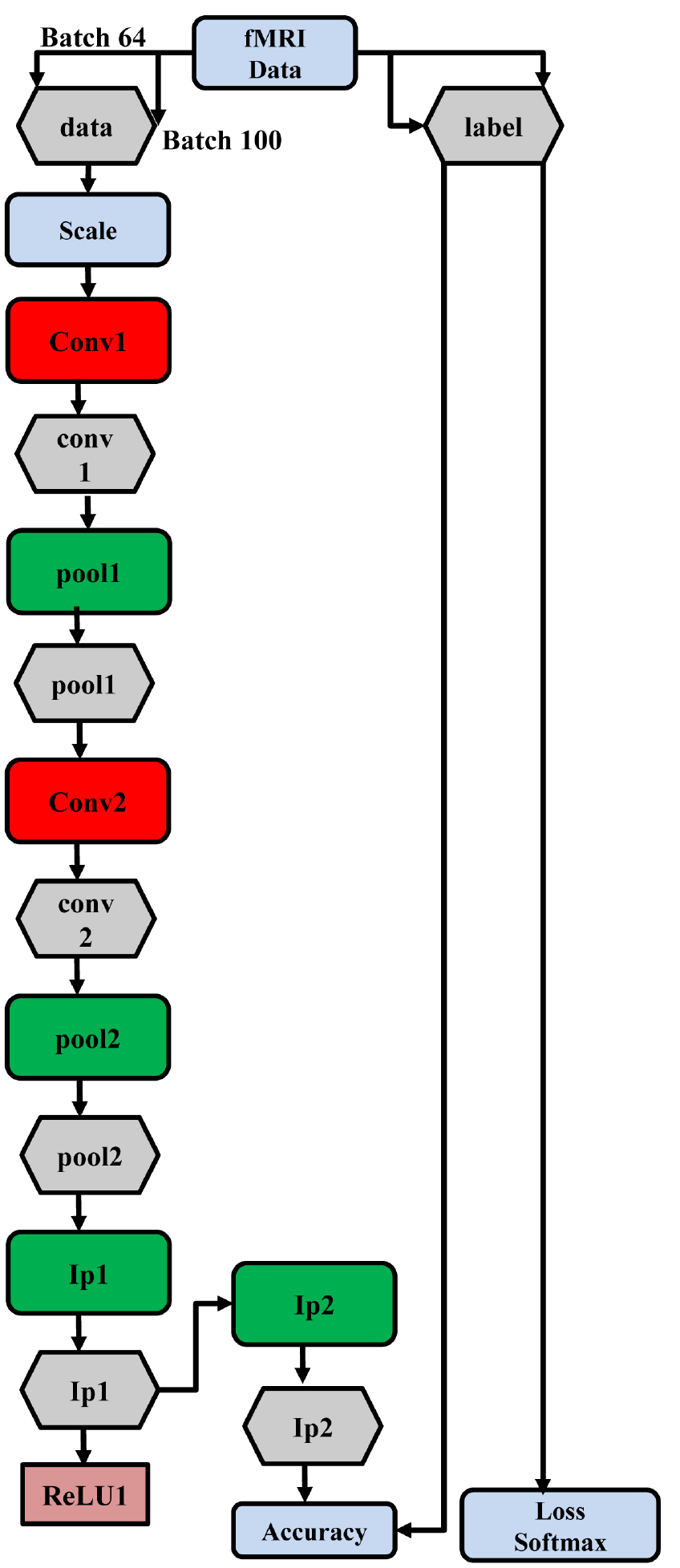
LeNet-5 network implemented for fMRI data

## III. Results and Discussion

The preprocessed fMRI 4D data in Nifti format was concatenated across z and t axes and then converted to a stack of 2D images in JPEG form using the neuroimaging packages Nibabel (http://nipy.org/nibabel/) and Python OpenCV (opencv.org). Next, images were labeled for binary classification of Alzheimers vs. normal data. The labeled images were converted to lmdb storage databases for high-throughput to be fed into a Deep Learning platform. The LeNet model which is based on Convolutional Neural Network architecture from Caffe DIGITS 0.2 - deep learning framework (Nvidia version) was used to perform binary image classification. The data were divided into three parts: training (60%), validation (20%), and testing (20%). The number of epochs was set to 30, and the batch size was 64, resulting in 126,990 iterations. The LeNet was trained by 270,900 samples and validated and tested by 90,300 images. In order to achieve the robustness and reproducibility of the deep neural network, the cross validation process was repeated five times (fivefold cross validation). This is presented in Table I. The mean of images was removed from data in the Deep Learning preprocessing step. In the training phase, loss of training and testing, as well as accuracy of the validation data, was measured. The learning rate fell dramatically following the 10th epoch and decreased slightly after the 20th epoch, as is shown in Figure 4. The Deep Learning LeNet model successfully recognized the Alzheimers data from the normal control data, and the average accuracy rate reached 96.8588%, as is depicted in Figure 3. The training and testing processes were performed on NVIDIA GPU Cloud Computing, which significantly improved the performance of the Deep Learning classifier.

**Fig. 3:**
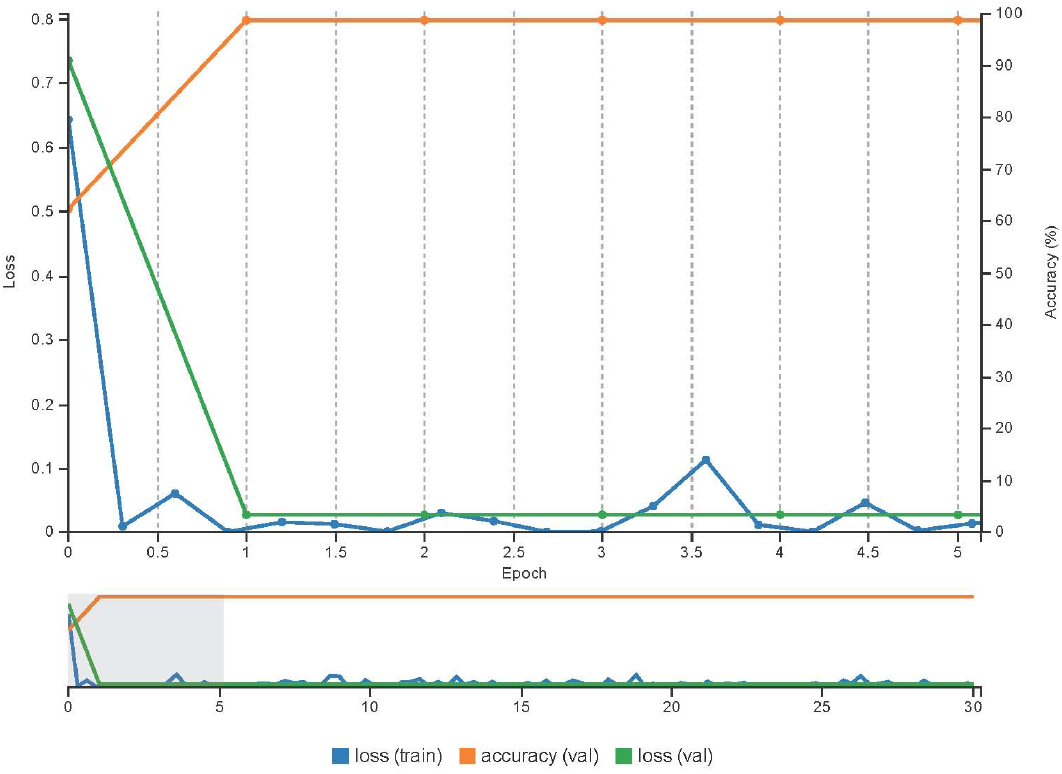
Loss train, Accuracy Validation (Test), Loss Validation for Run5

**Fig. 4:**
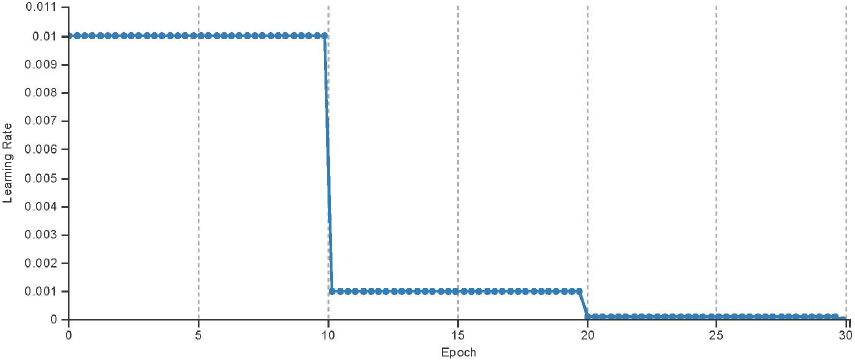
Learning Rate dropping each 10 epochs

**TABLE I:** Accuracies achieved from CNN across 5 runs

| Run1 | Run2 | Run3 | Run4 | Run5 | Mean |
| --- | --- | --- | --- | --- | --- |
| 96.858 | 96.857 | 96.854 | 96.863 | 96.8588 | **96.8588** |

Figure 4 demonstrates how the learning rate drops by 10 epochs during each of the training sets. We began with 0.01 and divided this by 10 for each of the 10 epochs.

Most challenges in traditional medical image processing and analyses have involved the selection of the best and most discriminative features, which must be extracted from data, and the process of choosing the best classification method. One important advantage of Deep Learning methods, especially the Convolutional Neural Network used in this study, is the ability to contain those two characteristics simultaneously. It is possible to visualize the results of filters (kernels) in each layer. Figures 5, 6 and 7 illustrate some of the final filter results from different layers. CNN is a strong feature extractor because of its convolutional layers, which are able to extract high-level features from images. This deep learning method is also a powerful classifier because of its complicated network architecture. The present solution, which is based on fully advanced preprocessing steps followed by CNN classification, improved the accuracy of AD data classification from 84% using Support Vector Machine (SVM) reported in the literature [14] [5] [12] to 96.86%. However, deep learning solutions have very few problems, such as high algorithm complexity and expensive infrastructure.

**Fig. 5:**
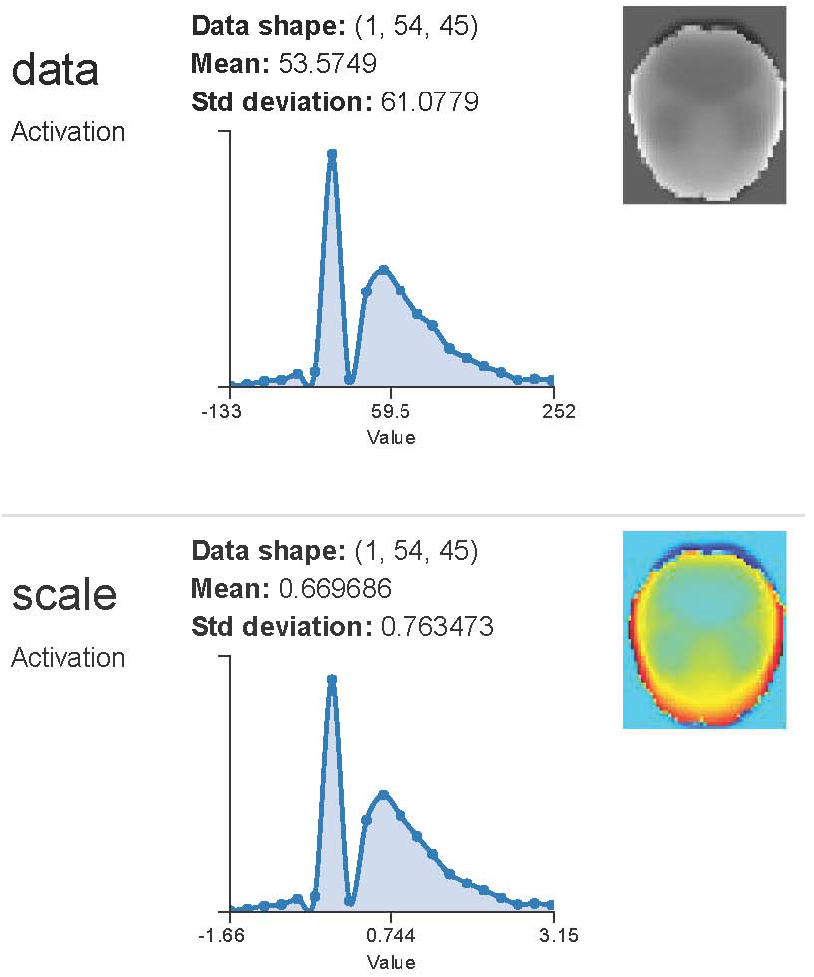
Statistics and Visualization for data and scale of Alzheimer’s Disease sample

**Fig. 6:**
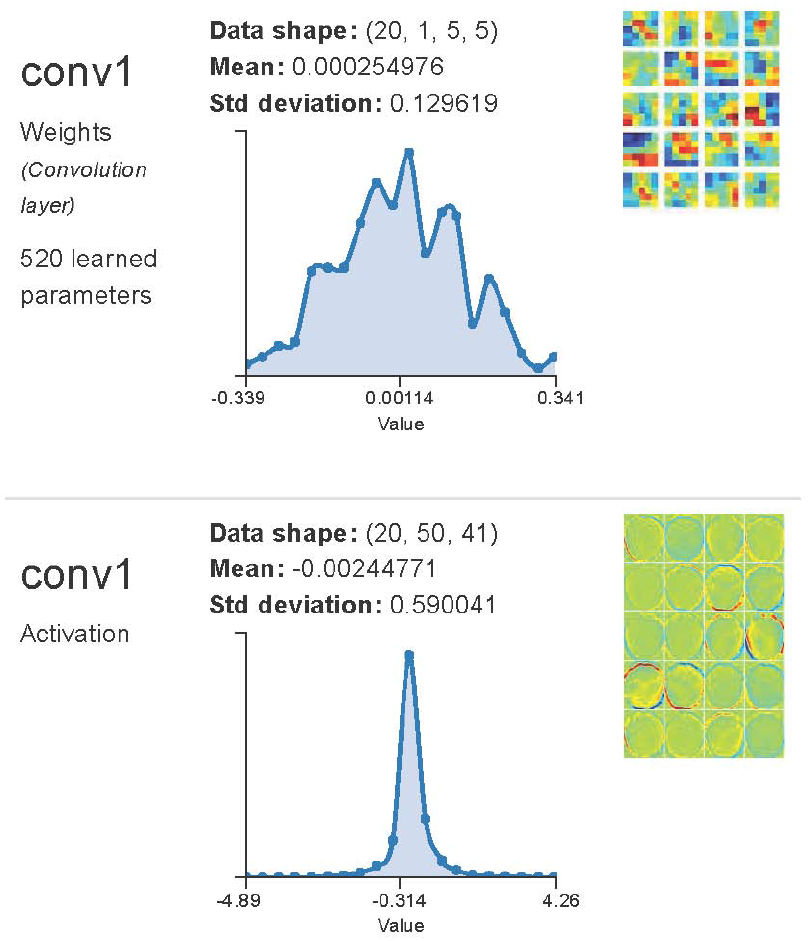
Statistics and visualization for first convolution layer of Alzheimer’s Disease sample

**Fig. 7:**
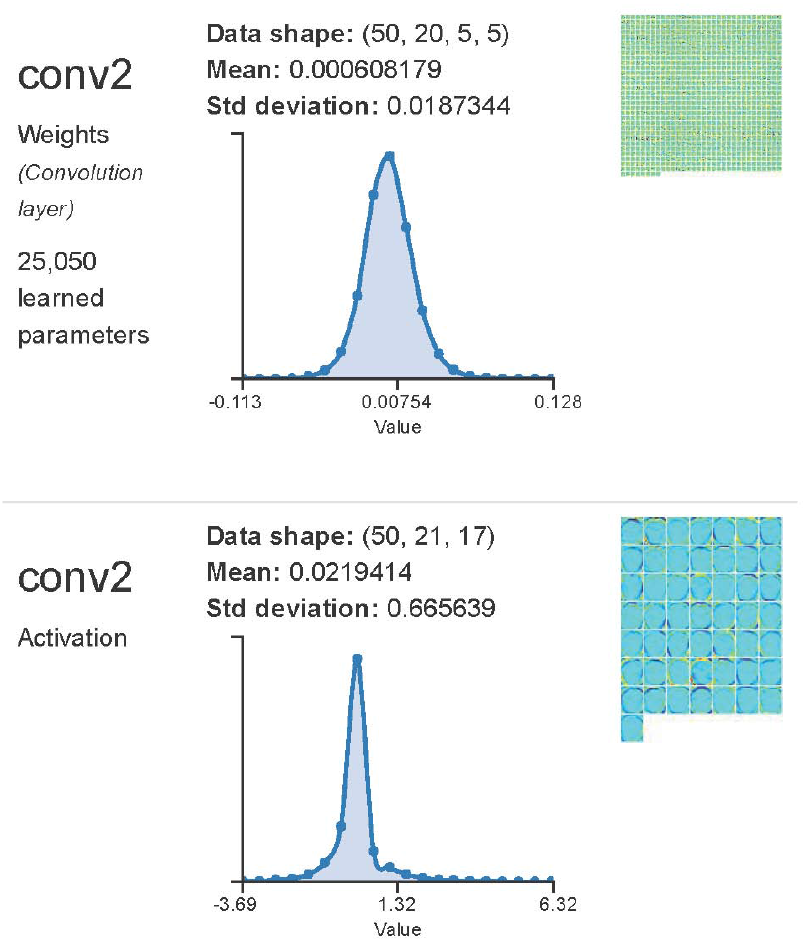
Statistics and visualization for second convolution layer of Alzheimer’s Disease sample

## IV. Conclusions

In this paper, we successfully classified AD data from normal control data with 96.86% accuracy using CNN deep learning architecture (LeNet), which was trained and tested with a massive number of images. This deep learning solution and the proposed pipeline not only open new avenues in medical image analyses, but also enable researchers and physicians to potentially predict any new data. It is also possible to generalize this method to predict different stages of Alzheimers disease for different age groups. Furthermore, this deep learning-based solution allows researchers to perform feature selection and classification with unique architecture. The rate of accuracy achieved in this work was very high, confirming that the network architecture was correctly selected. However, more complicated network architecture encompassing more convolutional neural layers is recommended for future work, and for more complicated problems.

## V. Acknowledgement

We would like to express our gratitude to Dr. Cristina Saverino, post-doctoral fellow at Toronto Rehabilitation Institute-University Health Network and Department of Psychology - University of Toronto, for extending her help and support throughout this study.

## VI. Appendix I

Data collection and sharing for this project was funded by the Alzheimer’s Disease Neuroimaging Initiative (ADNI) (National Institutes of Health Grant U01 AG024904) and DOD ADNI (Department of Defense award number W81XWH-12-2-0012). ADNI is funded by the National Institute on Aging, the National Institute of Biomedical Imaging and Bioengineering, and through generous contributions from the following: AbbVie, Alzheimer?s Association; Alzheimer’s Drug Discovery Foundation; Araclon Biotech; BioClinica, Inc.; Biogen; Bristol-Myers Squibb Company; CereSpir, Inc.; Eisai Inc.; Elan Pharmaceuticals, Inc.; Eli Lilly and Company; EuroImmun; F. Hoffmann-La Roche Ltd and its affiliated company Genentech, Inc.; Fujirebio; GE Healthcare; IXICO Ltd.; Janssen Alzheimer Immunotherapy Research & Development, LLC.; Johnson & Johnson Pharmaceutical Research & Development LLC.; Lumosity; Lundbeck; Merck & Co., Inc.; Meso Scale Diagnostics, LLC.; NeuroRx Research; Neurotrack Technologies; Novartis Pharmaceuticals Corporation; Pfizer Inc.; Piramal Imaging; Servier; Takeda Pharmaceutical Company; and Transition Therapeutics. The Canadian

Institutes of Health Research is providing funds to support ADNI clinical sites in Canada. Private sector contributions are facilitated by the Foundation for the National Institutes of Health (www.fnih.org). The grantee organization is the Northern California Institute for Research and Education, and the study is coordinated by the Alzheimer’s Disease Cooperative Study at the University of California, San Diego. ADNI data are disseminated by the Laboratory for Neuro Imaging at the University of Southern California.

## Footnotes

1 *Data used in preparation of this article were obtained from the Alzheimer’s Disease Neuroimaging Initiative (ADNI) database (adni.loni.usc.edu). As such, the investigators within the ADNI contributed to the design and implementation of ADNI and/or provided data but did not participate in analysis or writing of this report. A complete listing of ADNI investigators can be found at: http://adni.loni.usc.edu/wp-content/uploads/how_to_apply/ADNI_Acknowledgement_List.pdf

## References

[1] I. Arel, D. C. Rose, and T. P. Karnowski, “Deep machine learning-a new frontier in artificial intelligence research [research frontier],” Computational Intelligence Magazine, IEEE, vol. 5, no. 4, pp. 13–18, 2010.

[2] C. Grady, S. Sarraf, C. Saverino, and K. Campbell, “Age differences in the functional interactions among the default, frontoparietal control and dorsal attention networks,” Neurobiology of Aging, 2016.

[3] Y. Jia, E. Shelhamer, J. Donahue, S. Karayev, J. Long, R. Girshick, S. Guadarrama, and T. Darrell, “Caffe: Convolutional architecture for fast feature embedding,” in Proceedings of the ACM International Conference on Multimedia, pp. 675–678, ACM, 2014.

[4] Y. LeCun, L. Bottou, Y. Bengio, and P. Haffner, “Gradient-based learning applied to document recognition,” Proceedings of the IEEE, vol. 86, no. 11, pp. 2278–2324, 1998.

[5] A. Raventós and M. Zaidi, “Automating neurological disease diagnosis using structural mr brain scan features,”

[6] S. Sarraf, C. Saverino, H. Ghaderi, and J. Anderson, “Brain network extraction from probabilistic ica using functional magnetic resonance images and advanced template matching techniques,” in Electrical and Computer Engineering (CCECE), 2014 IEEE 27th Canadian Conference on, pp. 1–6, IEEE, 2014.

[7] S. Sarraf, E. Marzbanrad, and H. Mobedi, “Mathematical modeling for predicting betamethasone profile and burst release from in situ forming systems based on plga,” in Electrical and Computer Engineering (CCECE), 2014 IEEE 27th Canadian Conference on, pp. 1–6, IEEE, 2014.

[8] S. Sarraf and J. Sun, “Functional brain imaging: A comprehensive survey,” arXiv preprint arXiv:1602.02225, 2016.

[9] S. Sarraf and A. M. Golestani, “A robust and adaptive decision-making algorithm for detecting brain networks using functional mri within the spatial and frequency domain,” in The IEEE International Conference on Biomedical and Health Informatics (BHI), pp. 1–6, IEEE, 2016.

[10] S. M. Smith, M. Jenkinson, M. W. Woolrich, C. F. Beckmann, T. E. Behrens, H. Johansen-Berg, P. R. Bannister, M. De Luca, I. Drobnjak, D. E. Flitney, et al., “Advances in functional and structural mr image analysis and implementation as fsl,” Neuroimage, vol. 23, pp. S208–S219, 2004.

[11] S. C. Strother, S. Sarraf, and C. Grady, “A hierarchy of cognitive brain networks revealed by multivariate performance metrics,” in Signals, Systems and Computers, 2014 48th Asilomar Conference on, pp. 603–607, IEEE, 2014.

[12] E. E. Tripoliti, D. I. Fotiadis, and M. Argyropoulou, “A supervised method to assist the diagnosis and classification of the status of alzheimer’s disease using data from an fmri experiment,” in Engineering in Medicine and Biology Society, 2008. EMBS 2008. 30th Annual International Conference of the IEEE, pp. 4419–4422, IEEE, 2008.

[13] P. Vemuri, D. T. Jones, and C. R. Jack Jr, “Resting state functional mri in alzheimer’s disease,” Alzheimer’s research & therapy, vol. 4, no. 1, pp. 1–9, 2012.

[14] X. Zhang, B. Hu, X. Ma, and L. Xu, “Resting-state whole-brain functional connectivity networks for mci classification using l2-regularized logistic regression,” NanoBioscience, IEEE Transactions on, vol. 14, no. 2, pp. 237–247, 2015.

